# Impact of nephrotoxins and oxidants on survival and transport function of hiPSC-derived renal proximal tubular cells

**DOI:** 10.1101/2025.02.17.638605

**Authors:** Isaac Musong Mboni-Johnston, Sören Hartmann, Christian Kroll, Carsten Berndt, James Adjaye, Nicole Schupp

## Abstract

Due to their role in excretion, renal proximal tubule cells are susceptible to damage by toxic metabolites and xenobiotics. The regenerative capacity of the kidney allows for the replacement of damaged cells, a process involving differentiation programs. However, kidney function tends to decline, suggesting that the replacement cells may not achieve full functionality. To understand possible causes of this decline, we investigated effects of nephrotoxins and oxidants on the differentiation of induced pluripotent stem cells (iPSC) into proximal tubular epithelial-like cells (PTELC). Proliferation, apoptosis, senescence and expression of oxidative defense genes were analyzed in iPSCs, differentiating and differentiated cells treated with cisplatin (CisPt, up to 45 µM), cyclosporin A (CycA, up to 12 µM) and the oxidants menadione (Mena, up to 50 µM) and tert-butylhydroquinone (tBHQ, up to 50 µM). We found that differentiating cells were most sensitive to oxidants and showed increased sensitivity to CisPt, whereas all differentiation stages showed similar sensitivity to CycA. Both oxidative stress and CisPt triggered apoptosis in all differentiation stages, whereas CycA mainly induced senescence. Treatment during differentiation resulted in long-term effects on gene expression in differentiated cells. While oxidants had no effect on transport function of differentiated cells, CisPt and CycA impaired albumin uptake. Our data suggest a substantial sensitivity of differentiating cells to nephrotoxins and oxidants, an aspect that could potentially interfere with regenerative processes.

## Introduction

As an important route for excretion of drugs and other chemicals from the body, the kidneys play a crucial role in maintaining homeostasis. However, this essential function also makes the kidney cells very susceptible to damage from xenobiotics and their toxic metabolites (Choudhury et al. 2017; Tiong et al. 2014). In particular, the epithelial cells lining the proximal tubule are affected by toxic substances due to their role in glomerular filtrate concentration, transport and metabolism of compounds that can trigger acute kidney injury (AKI). Therefore, proximal tubular epithelial cells (PTEC) have become the focus of extensive research in renal toxicity assessment and regenerative medicine (Naughton 2008; Szeto and Chow 2005).

To ensure the organ’s functionality, cells must be replaced constantly, even if there is no specific injury, as the cells age. These regenerative processes reach a higher level in the case of kidney damage compared to basal cell turnover (Ameku et al. 2016; Castrop 2019). Tubular regeneration is achieved either by a local stem cell population or by the dedifferentiation of surviving PTEC which allows them to regain proliferation. Either one or an interplay of both regenerative pathways maintains the organ’s functionality and structure (Berger and Moeller 2014; Bonventre 2003). However, despite the regenerative capacity of PTEC in the kidney, the regeneration processes are often fraught with challenges. If they are inefficient, impaired and dysregulated, some replaced cells may not regain full functionality after injury (Venkatachalam et al. 2018; Venkatachalam et al. 2015). In addition, AKI patients may have impaired kidney function even after recovery, increasing their risk of developing chronic kidney disease (CKD) and end-stage renal disease (ESRD). This could be a consequence of not all damaged cells having been replaced by fully functional cells or their functionality having been impaired during the regeneration process (Coca et al. 2012; Goldstein et al. 2013; Rangaswamy and Sud 2018; Venkatachalam et al. 2018).

Since ineffective regeneration is linked to the progression of AKI to CKD and ESRD, it is essential to understand the causes of this reduced functionality of tubular epithelial cells after regeneration. One common cause of impaired renal function observed in clinical practice is drug induced nephrotoxicity. Drugs in clinical use often trigger off-target effects on PTEC which have been implicated in 19-25 % of AKI in critically ill patients (Mehta et al. 2004). In addition to the non-target effects of drugs on PTEC, reactive oxygen species (ROS) and oxidative damage have been implicated to be common and significant pathobiological factors in PTEC injury (Bhargava and Schnellmann 2017). While ROS play an essential role in intracellular signaling and cell homeostasis, excessive ROS cause oxidative damage to all biomolecules. PTEC are a significant source of ROS in the kidney due to their high mitochondrial density, which is necessary for their enormous energy requirements to maintain secretory and absorptive functions (Bhargava and Schnellmann 2017). Consequently, PTEC are highly vulnerable to ROS-induced damage leading to AKI (Percy et al. 2008). Given the role of nephrotoxic drugs and oxidative stress in PTEC injury, these stressors can be expected to compromise the integrity and functionality of newly replaced PTEC during regeneration and may lead to maladaptive repair. Therefore, studies are needed that explore the effects of nephrotoxins and oxidative stress on renal regeneration and thus go beyond the mere prediction of nephrotoxicity on mature PTEC.

A promising strategy to investigate this is the use of human induced pluripotent stem cell (hiPSC)-based models (Zink et al. 2020). As proof of principle for the use of hiPSCs to model renal regeneration, using a protocol previously reported by Kandasamy et al. (Kandasamy et al. 2015), we recently showed a hiPSC-based *in vitro* model that allows toxicity assessment during PTEC differentiation from hiPSCs (Mboni-Johnston et al. 2023). The results showed that hiPSC differentiation resulted in cells that resembled PTECs not only morphologically but also in their mRNA and protein expression patterns and functionality. When treating the cells in different differentiation stages with known nephrotoxins, we also observed that the cells in the differentiation process reacted more sensitively to the toxins than the fully differentiated cells. This increased sensitivity of the differentiating cells could have an unfavorable effect on regeneration processes (Mboni-Johnston et al. 2023). Here, we not only characterized cellular mechanisms that might contribute to the sensitivity of differentiating cells to nephrotoxins but also investigated the effects of oxidative stress-inducing substances which may be present in the kidney during AKI.

## Methods

The authors declare that all supporting data are available within the article and its online-only Data Supplement.

### Materials

The human induced pluripotent stem cell line Foreskin-4 (iPS(foreskin) clone (#4), WB66699), derived from human foreskin fibroblasts by lentiviral transfection of OCT3/4, SOX2, NANOG, and LIN28 (Yu et al. 2007), from now on referred to as hiPSC, was purchased from WiCell at passage 27 and used until passage 47 (Madison, USA). Human embryonic stem cell qualified Matrigel and Reduced Growth Factor Basement Membrane Matrix were from Corning (New York, USA), mTeSR1 medium from StemCell Technologies (Vancouver, Canada), and renal epithelial growth medium (REGM) containing various growth factors and supplements (REGM BulletKit) from Lonza (Basel, Switzerland). Y-27632 dihydrochloride and BMP-2 were from Sigma-Aldrich (St. Louis, USA), and BMP-7 was supplied by Thermo Fisher Scientific (Waltham, USA). All chemicals were purchased from Sigma-Aldrich (St. Louis, USA), Enzo Life Sciences (Farmingdale, NY, USA), and Teva (Petach Tikva, Israel). Primers (Table S1) were from Eurofins Genomics (Val Fleuri, Luxembourg), antibodies (Table S2) were from Abcam (Cambridge, UK) and BD Bioscience (Heidelberg, Germany), and the fluorophore-conjugated secondary antibody Alexa Fluor 488 was from Life Technologies (Carlsbad, California, USA).

### Cultivation of hiPSC and differentiation procedure

hiPSC were cultured on 6-well plates plated with human embryonic stem cell-qualified Matrigel in serum-free, defined mTeSR1 medium supplemented with 10 mM Y-27632 dihydrochloride. Authentication was done before purchasing and after completion of experiments. In addition to this, the cells were also regularly checked for mycoplasma, bacteria, and fungi. The cells were maintained according to WiCell’s recommendations. This included thawing a vial in three wells of a 6-well plate and culturing in a humidified atmosphere with 5 % CO_2_ at 37 °C as well as twice weekly passaging at a ratio of 1:6 when the cultures had reached 70-90 % confluence to keep them in exponential growth. A modified protocol, according to Kandasamy and colleagues (Kandasamy et al. 2015), was used for differentiation into renal PTELC, as previously reported (Mboni-Johnston et al. 2023).

### Analysis of cell viability and antioxidative capacity

Undifferentiated hiPSC, hiPSC differentiating into PTELC, and hiPSC differentiated into PTELC were treated with distinct concentrations of CisPt, CycA, Mena, and tBHQ. After 24 h of exposure, the viability of the undifferentiated, differentiating, and differentiated cells was examined using the Alamar Blue Assay (O’Brien et al. 2000), as described previously (Mboni-Johnston et al. 2023). The relative viability of the cells in the corresponding untreated controls was set to 100 %.

GSH and GSSG were measured with the GSH/GSSG-Glo assay (Promega, Walldorf, Germany) according to the manufacturer’s instructions.

### Analysis of gene expression (RT-PCR)

RNA was isolated using the RNeasy Mini Kit and RNase-free DNase Set (Qiagen, Venlo, The Netherlands) according to the manufacturer’s protocol as reported previously (Mboni-Johnston et al. 2023). The primer sequences (Table S1, supplement) used for the mRNA expression analyses were designed with the help of NCBI (https://www.ncbi.nlm.nih.gov/nuccore/) and validated for specificity. At the end of the run, data were analyzed using CFX Manager software (v3.1, Bio-Rad Laboratories, Hercules, CA, USA) according to the manufacturer’s instructions. All samples were run with three technical replicates. The Cq values obtained for the genes of interest were first normalized to the mean Cq values obtained for the housekeeping genes ACT-β and RPL-32 and then for the control samples and expressed as a fold of this mean value. Changes in gene expression of ≤0.5- and ≥2- fold were considered biologically relevant.

### Analysis of cell proliferation, apoptosis, and function

To investigate the effect of toxins on the proliferation rate, the incorporation of EdU into S-phase cells was determined using the EdU-Click 488 assay according to the manufacturer’s protocol (Baseclick GmbH, Tutzing, Germany) as reported earlier (Mboni-Johnston et al. 2023).

To verify whether the substances analyzed induce apoptosis in the cells at different stages of differentiation, the TUNEL (terminal deoxynucleotidyl transferase-mediated dUTP-fluorescein nick-end labeling) assay was performed with cells grown on coverslips using the In-Situ Cell Death Detection Assay Kit (Roche, GmbH, Germany) according to the manufactureŕs instructions. The TUNEL assay visualizes free hydroxyl groups from fragmented DNA produced during apoptosis by labeling them with modified nucleotides through an enzymatic reaction (Gavrieli et al. 1992). The coverslips were mounted with VECTASHIELD® Antifade Mounting Medium containing DAPI and sealed with nail polish. Fluorescence images were acquired using a fluorescence microscope (Olympus BX43 Upright Microscope, Olympus, Shinjuku, Tokyo, Japan) at 200-fold magnification and analyzed using ImageJ 1.51j8 (Schneider et al. 2012). TUNEL-positive cells were referenced to the total number of cells and normalized to the untreated control.

The effect of toxins on the functionality of hiPSC-derived PTELC was examined by adding fluorescently labeled bovine serum albumin (FITC-BSA) (Sigma Aldrich, 10 mg/ml) to the culture medium. Cells were incubated with serum-free medium with/without 100 µg/ml FITC-albumin for 2 h at 37°C, and uptake was stopped with ice-cold PBS. Cells were then harvested from Matrigel-coated wells with trypsin/EDTA, and albumin uptake was analyzed by flow cytometry (Becton Dickinson, Accuri™ C6 plus (Heidelberg, Germany)) as previously reported (Mboni-Johnston et al. 2023).

### Statistical Analysis

Statistical analysis was performed using GraphPad Prism version 6 (GraphPad Software, San Diego, CA, USA). Data are presented as the mean with standard deviation of three independent experiments (n = 3), RT-qPCR results also included 3 technical replicates. Normal distribution was checked using the Kolmogorov-Smirnov test. For comparisons of two normally distributed groups the two-tailed unpaired Student’s t-test and for non-normally distributed values, the Mann-Whitney U-test was used. Multiple groups were tested by one-way analysis of variance (one-way ANOVA) with Tukey’s or Dunnett’s post hoc test. Non-parametrical datasets of multiple groups were analyzed with Kruskal-Wallis one-way ANOVA on ranks test. Statistically significant differences between the groups were assumed at a p-value ≤0.05.

### Results

An *in vitro* model based on human induced pluripotent stem cells (hiPSC, Figure 1A) was used to investigate the influence of the nephrotoxins cisplatin (CisPt) and cyclosporin A (CycA) as well as the oxidizing agents menadione (Mena) and tert-butylhydroquinone (tBHQ) on renal tubular differentiation processes. Treatment with the respective compounds was performed on undifferentiated hiPSC, differentiating cells at day three of differentiation (diffD3) and differentiated cells after nine days of differentiation (diffD9). Analyses were performed after 24 h of exposure, as indicated in Figure 1A. First, IC_20_ and IC_50_ (inhibitory concentrations where 20 and 50 % of the cells are dead, respectively) needed to be determined for the oxidants, as shown in Figures 1B and C. As we have already investigated the sensitivity of the cells at different stages of differentiation to the nephrotoxins CisPt and CycA (Mboni-Johnston et al. 2023), the respective IC_20_ and IC_50_ values determined in this former study were used here for further investigations.

**Figure 1.**
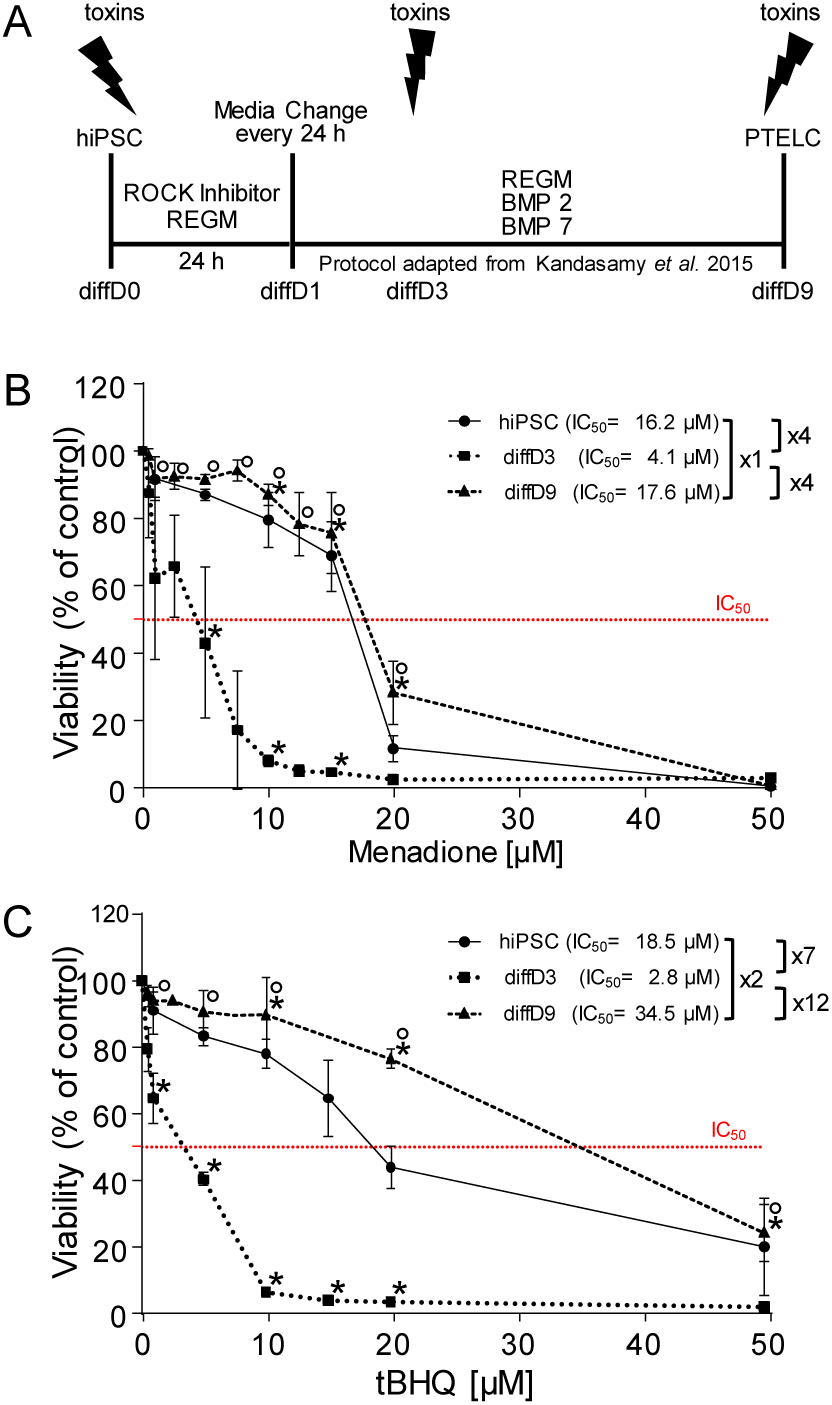
Influence of selected oxidizing agents on the viability of hiPSC and cells at differentiation days 3 and 9. (A) Schematic representation of the protocol for differentiating hiPSC into PTELC showing the treatment time points used during the differentiation process. (B and C) Cell viability measured using the Alamar Blue assay on hiPSC and cells treated at differentiation days 3 (diffD3), and 9 (diffD9) for 24 hours with the indicated concentrations of (B) menadione and (C) tert-butylhydroquinone (tBHQ). The red dotted line indicates 50 % viability and therefore the IC_50_. Data from at least 3 independent experiments are given as mean ± SD. *p≤0.05 vs. hiPSC, °p≤0.05 vs. diffD3 (one-way ANOVA). BMP = bone morphogenetic protein, diffD = differentiation day, hiPSC = human induced pluripotent stem cells, IC_50_ = inhibitory concentration where 50 % of the cells are dead, PTELC = proximal tubular epithelial-like cells, REGM = renal epithelial cell growth medium, ROCK = rho-associated, coiled-coil-containing protein kinase.

#### hiPSC-derived tubular progenitors are particularly sensitive to oxidative stress

hiPSC, diffD3, and diffD9 were treated with two different oxidants, Mena and tBHQ, for 24 h, and their sensitivity to the respective substances was analyzed using the Alamar Blue assay. diffD3 showed the highest sensitivity to Mena, with an IC_50_ about four times lower than that of hiPSC and diffD9, which had a very similar IC_50_ (Figure 1B). When exposed to the second oxidizing agent, tBHQ, diffD3 also showed the highest sensitivity with an IC_50_ 7- to 12-fold lower than those of hiPSC and diffD9 (Figure 1C). Here, hiPSC were twice as sensitive to tBHQ compared to diffD9, which showed the highest resistance. Overall, the data demonstrate that cells with the same genetic background exhibited differential sensitivity to oxidants at different differentiation stages, with hiPSC differentiating into PTELC being the most sensitive.

#### Impact of differentiation on the antioxidative defense

To elucidate why diffD3 cells were so sensitive to treatment with oxidants, mRNA expression of genes associated with antioxidative defense was quantified. When comparing the expression of selected genes with the expression in hiPSC, diffD9 showed a significant increase in four of seven tested genes, while only glutathione peroxidase (Gpx1) was significantly upregulated in diffD3 (Figure 2A). Five of the seven tested genes were significantly expressed at lower levels in diffD3 than in diffD9, which was then tested on a larger set of antioxidative defense genes. Of the fourteen tested genes, eight were significantly reduced in diffD3 compared to diffD9 and only two, a pro-oxidative gene, the subunit of NADPH oxidase (p47) and the master regulator of the antioxidative defense, nuclear factor erythroid 2-related factor 2 (Nrf2), were significantly upregulated (Figure 2B). Antioxidant capacity, measured as the ratio of reduced glutathione to oxidized glutathione, was significantly higher in diffD9 compared to hiPSC, while the high variance of diffD3 results did not lead to a significant result (Figure 2C). Quantification of the expression of genes responsible for the synthesis of glutathione revealed a significant downregulation of both, glutamate-cysteine ligase (Gclc) and glutathione synthetase (Gss) in diffD3 as well as in diffD9 (Figure 2D).

**Figure 2:**
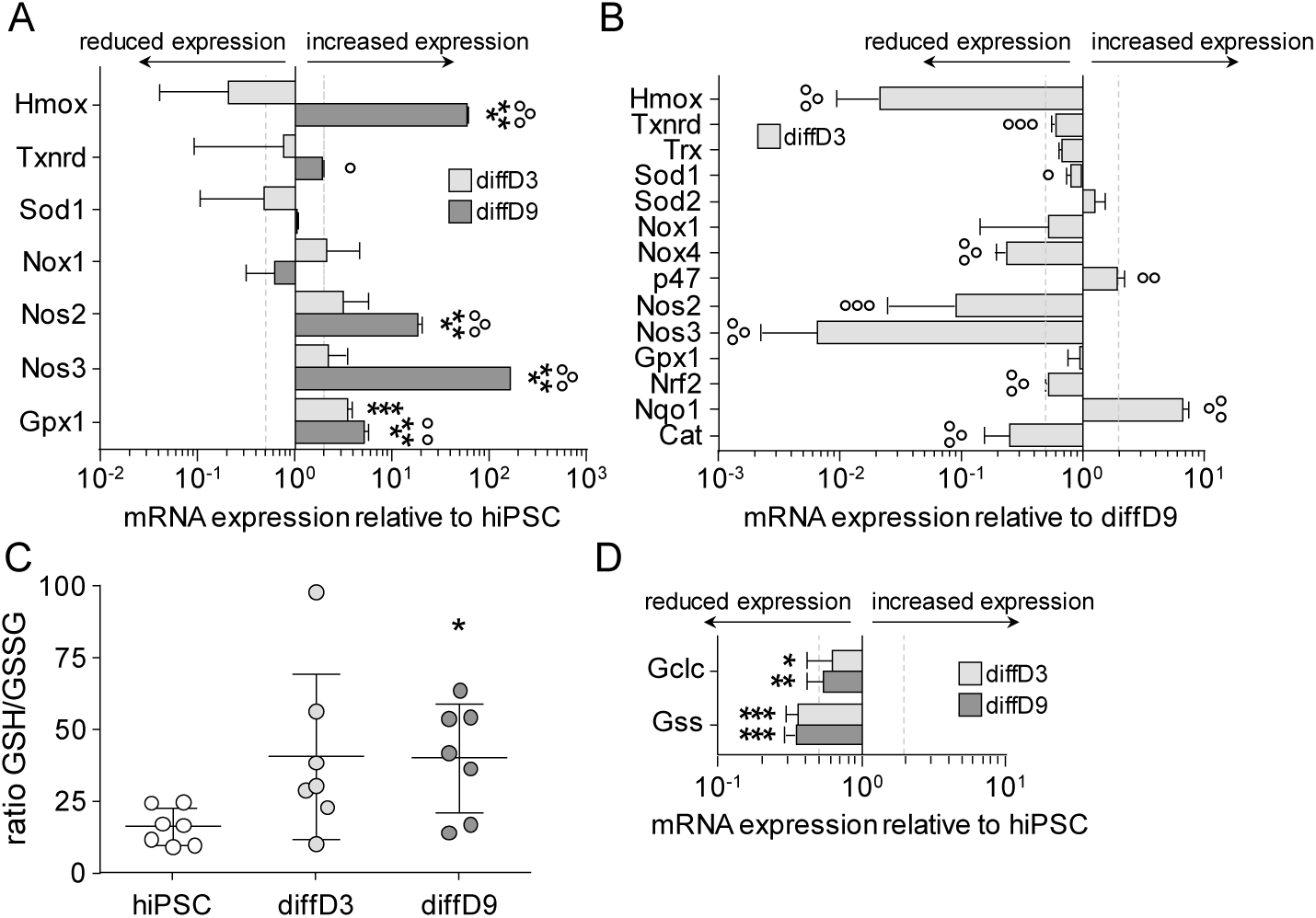
Expression of antioxidative defense genes and antioxidative capacity of differentiating and differentiated cells. (A) mRNA expression of antioxidative defense genes in diffD3 and diffD9 compared to hiPSC. (B) mRNA expression of antioxidative defense genes in diffD3 compared to diffD9. (C) Measurement of the antioxidative capacity of hiPSC, diffD3 and diffD9 as the ratio of reduced to oxidized glutathione. (D) mRNA expression of glutathione synthesis genes in diffD3 and diffD9 compared to iPSC. Data from at least 3 independent experiments are expressed as mean + SD. *p≤0.05, **p<0.01, and ***p<0.001 compared to hiPSC (One-way ANOVA (A,C) or Student’s t-test (D)), ° p≤0.05, °°p<0.01, °°°p<0.001 vs. diffD3 (A,C, One-way ANOVA) or diffD9 (B, Student’s t-test). Cat = catalase, diffD3/9 = differentiation day 3/9, hiPSC = human induced pluripotent cells, Hmox = heme oxygenase-1, Gclc = glutamate-cysteine ligase, catalytic subunit, Gpx1 = glutathione peroxidase 1, Gss = glutathione synthetase, Nox1/4 = NADPH oxidase catalytic unit 1/4, Nos2/3 = nitric oxide synthase 2/3, Nqo1 = NAD(P)H quinone dehydrogenase 1, Nrf2 = nuclear factor erythroid 2-related factor 2, p47 = p47phox, a subunit of NADPH oxidase, Sod1/2 = superoxide dismutase 1/2, Trx = thioredoxin, Txrnd = thioredoxin reductase 1.

#### Influence of oxidizing agents on gene expression and transport in proximal tubular epithelial-like cells

In addition to the disruptive impact of the nephrotoxins CisPt and CycA on gene expression and function of differentiated PTELC, reported before (Mboni-Johnston et al. 2023), here, we have analyzed the effect of the oxidants Mena and tBHQ. Our results show that treatment of diffD9 with Mena selectively affected mRNA expression of individual PTEC markers and transporters (Figure 3A). There was a significant and dose-dependent decrease in the mRNA expression of aquaporin 1 (Aqp-1) and megalin (Meg) in cells treated with both IC_20_ and IC_50_, as well as a reduced expression of peptide transporter (Pept-) 2 observed after treatment with the IC_50_. Alanyl aminopeptidase (CD13) and Pept-1 were the only genes that increased significantly after treatment with IC_50_ of Mena, almost reaching a 2-fold amount. The changes in the other genes were nowhere near the 0.5- or 2-fold change considered biologically relevant. When treated with tBHQ (Figure 3B), of the PTEC-specific genes analyzed, Aqp-1 was the only gene that showed a significant dose-dependent decrease after treatment with IC_20_ and IC_50_. mRNA expression levels of cadherin (Cad-) 16, Meg, and Pept-1 increased above 2-fold after treatment with the IC_50_, with only Cad-16 reaching statistical significance. mRNA expression of the pluripotency marker Oct-3/4 showed no significant change while Nanog, was significantly increased by the IC_20_ of Mena.

**Figure 3:**
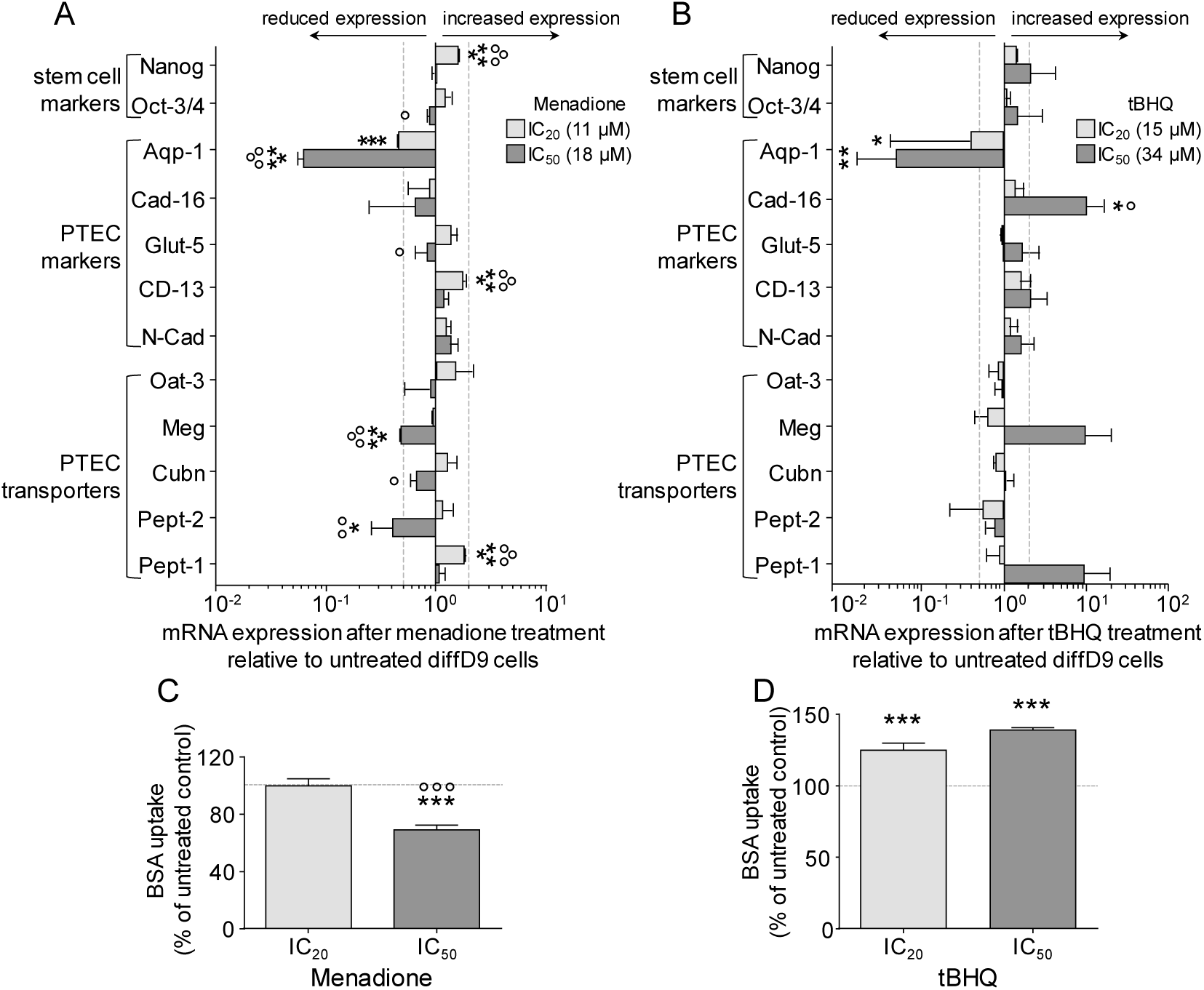
Influence of oxidizing agents on marker gene expression and transport function in hiPSC differentiated into proximal tubular epithelial cells. mRNA expression of marker genes in differentiated (diffD9) cells after 24-hour treatment with the IC_20_ and IC_50_ concentrations of (A) Mena and (B) tBHQ compared to their expression in untreated diffD9, analyzed by quantitative RT-PCR. Flow cytometric analysis of albumin uptake in diffD9 after 24 hours of treatment with the IC_20_ and IC_50_ levels of (C) Mena and (D) tBHQ using FITC-labelled albumin. Data from at least 3 independent experiments are expressed as mean + SD. *p≤0.05, **p<0.01, and ***p<0.001 compared to untreated diffD9 cells, ° p≤0.05, °°p<0.01, °°°p<0.001 vs. cells treated with IC_20_ (One-way ANOVA). Aqp-1 = aquaporin-1, Cad-16 = cadherin 16, CD13 = alanyl aminopeptidase, Cubn = cubilin, diffD9 = differentiation day 9, Glut-5 = fructose transporter 5, hiPSC = human induced pluripotent cells, IC_20_/_50_ = inhibitory concentration where 20 and 50 % of the cells are dead, respectively, Mena = menadione, Meg = megalin, Nanog = homeobox gene, Oat-3 = organic anion transporter 1, Oct-3/4 = octamer-binding transcription factor 3/4, Pept-1/2 = peptide transporter 1, PTEC = proximal tubular epithelial cells, tBHQ = tert-butylhydroquinone.

In analogy to the nephrotoxins CisPt and CycA, which reduced the albumin uptake capacity of PTELC (Mboni-Johnston et al. 2023), we also investigated the effects of the two oxidants on this cell function (Figure 3C and D). A significant decrease in albumin transport was observed after treatment with IC_50_ of menadione, while tBHQ increased the transport.

#### Investigation of the influence of nephrotoxins and oxidants on the fate of cells at distinct stages of differentiation

Once the sensitivity of the three differentiation stages to the various toxins was known, the effects of the toxins on proliferation, apoptosis, and senescence were investigated. To gain insight into the impact of the substances on proliferation, cells at the three differentiation stages were treated with the hiPSC’s IC_20_ and IC_50_ of CisPt, CycA, Mena and tBHQ. Cells in the S-phase were visualized with the help of the fluorescent base analogue 5-ethynyl-2′-deoxyuridine (EdU). Treatment with CisPt inhibited the likely proliferation-associated EdU incorporation in hiPSC in a dose-dependent manner (Figure 4A and B). However, this was not the case in diffD3 and diffD9, in which EdU incorporation was not affected. The proliferation of hiPSC treated with CycA decreased by approximately 30 % compared to the untreated control, but only significantly at the IC_50_ (Figure 4C and D). No significant difference in EdU incorporation was observed in CycA-treated diffD3 at both concentrations tested compared to the untreated controls, while the number of cells in S-phase was reduced by about half in diffD9 after treatment with CycA compared to the untreated control, although this was only significant at the IC_20_. Surprisingly, oxidants diminished proliferation significantly only in diffD3 compared to the untreated control. In contrast, no significant differences in EdU incorporation were observed when hiPSC and diffD9 were treated with Mena and tBHQ (Figures 3E to H).

**Figure 4:**
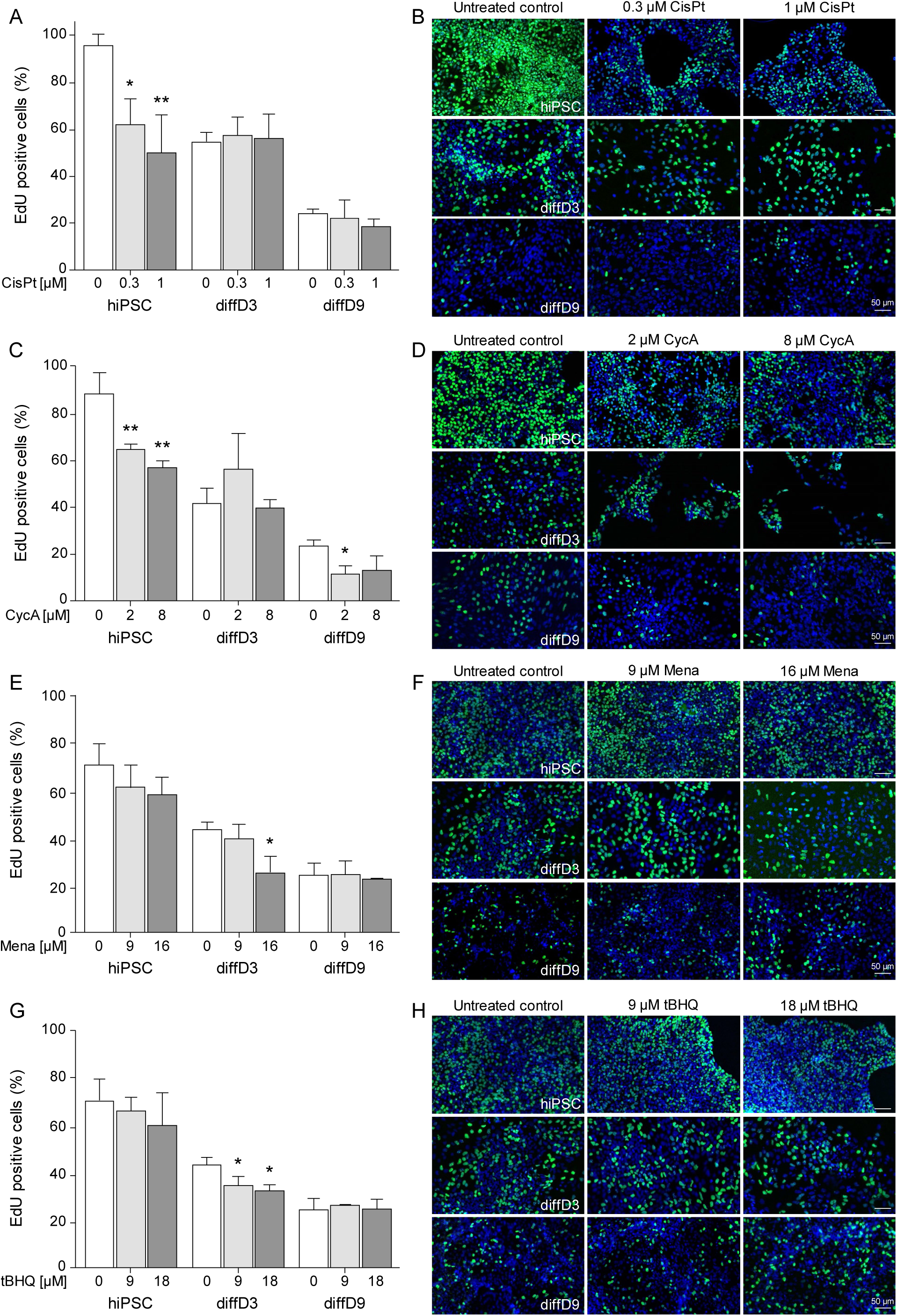
Influence of genotoxins and oxidizing agents on proliferation. hiPSC, diffD3, and diffD9 were treated for 24 h with the indicated concentrations of CisPt (A, B), CycA (C, D), Mena (E, F) and tBHQ (G, H) corresponding to the hiPSC’s IC_20_ and IC_50_ of the respective substances. After treatment, cell proliferation was determined by measuring the incorporation of EdU. (A, C, E, G) Quantification of EdU-positive cells from 7-10 visual fields. Data from at least three independent experiments are expressed as mean + SD. One-way ANOVA, *p≤0.05 and **p<0.01 compared to untreated control. (B, D, F, H) Representative merged images of EdU-positive cells (FITC, green) and cell nuclei (DAPI, blue) from treated and untreated hiPSC, diffD3, and diffD9 cells from three independent experiments are shown. The scale bar represents 50 µm. CisPt = cisplatin, CycA = cyclosporin A, DAPI = 4′,6-diamidino-2-phenylindole, diffD3/9 = differentiation day 3/9, EdU = 5-Ethynyl-2′-deoxyuridine, FITC = fluorescein isothiocyanate, hiPSC = human induced pluripotent cells, IC_20_/_50_ = inhibitory concentration where 20 and 50 % of the cells are dead, respectively, Mena = menadione, tBHQ = tert-butylhydroquinone.

The induction of apoptosis by the four substances was analyzed with the TUNEL assay. Cells of the three differentiation stages were treated with the respective equitoxic concentrations (IC_20_ and IC_50_ of the hiPSC) of CisPt, CycA, Mena, and tBHQ. In the case of CisPt, for the treatment of the diffD9 cells, a higher concentration was added, since the IC_50_ of the hiPSC did not lead to a significant number of apoptotic cells. As shown in Figure 5A, the frequency of TUNEL-positive cells increased in a dose-dependent manner after CisPt treatment in hiPSC, diffD3, and diffD9. When treated with CycA, the frequency of TUNEL-positive cell nuclei also increased with rising concentrations in hiPSCs, diffD3, and diffD9, but not as much as after CisPt treatment and reached significance only with the higher concentration in diffD3 and diffD9 (Figure 5B). In the case of the oxidizing agents, the frequency of TUNEL-positive cells was generally much lower than for the other two substances. It was significantly increased only in Mena- (Figure 5C) and tBHQ- (Figure 5D) treated diffD3.

**Figure 5:**
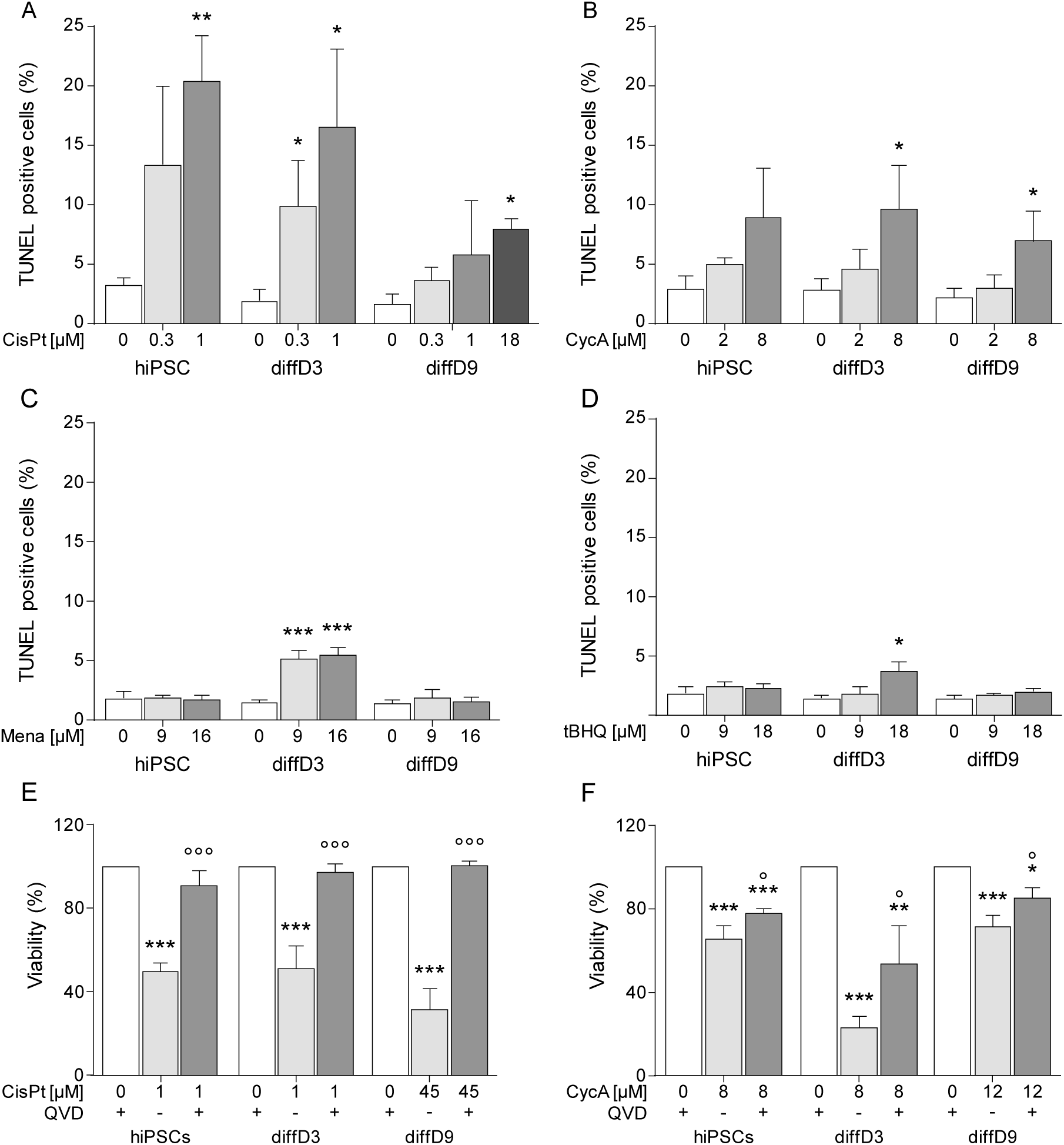
Stimulation of apoptosis in hiPSC, diffD3, and diffD9 by CisPt, CycA, Mena and tBHQ and verification by pan-caspase inhibition. hiPSC, diffD3, and diffD9 were treated with equitoxic doses of CisPt (A), CycA (B), Mena (C) and tBHQ (D) for 24 hours. After treatment, apoptosis was measured with the TUNEL assay and quantified by counting the number of TUNEL-positive cells. Cell viability of cells of all differentiation stages after treatment with CisPt (E) and CycA (F) with and without the addition of 10 µM of the pan-caspase inhibitor QVD. All data are presented as mean + SD of three independent experiments. One-way ANOVA, *p≤0.05, **p<0.01, ***p<0.001 vs. the respective untreated control and °p ≤0.05 and °°°p ≤0.001 compared to treated cells without QVD in E and F. CisPt = cisplatin, CycA = cyclosporin A, diffD3/9 = differentiation day 3/9, hiPSC = human induced pluripotent cells, Mena = menadione, QVD = quinoline-Val-Asp-difluorophenoxymethylketone, tBHQ = tert-butylhydroquinone, TUNEL = terminal deoxynucleotidyl transferase-mediated dUTP (deoxyuridine triphosphate)-fluorescein nick-end labelling.

A pan-caspase inhibitor was used to determine whether the cell death induced by the two substances CisPt and CycA, which triggered a relevant amount of apoptosis compared to the oxidants, was predominantly due to this apoptosis. Cell viability data showed significant restoration of cell viability to almost control level in all stages of differentiation under co-treatment with CisPt and the pan-caspase inhibitor QVD compared to the CisPt mono-treatment (Figure 5E). Also, the percentage of viable cells at the different stages of differentiation treated with CycA and QVD together was significantly higher than when treated with CycA alone (Figure 5F). However, in contrast to CisPt, the viability of the cells could not be rescued entirely.

Since all substances except CisPt did not impair cell viability primarily via apoptosis, the triggering of senescence by the toxins was investigated. As expected, 48 h after a 24 h treatment with CisPt, no extensive expression of the group of genes linked to the senescence-associated secretory phenotype was observed in cells of different stages of differentiation; only p21 was significantly higher expressed in diffD3 (Figure 6A). CycA, on the other hand, stimulated significant expression of the senescence-associated genes IL-8, p21, as well as β-GAL in hiPSC (Figure 6B). In diffD3, no gene was significantly or relevantly increased, while in diffD9, both IL-8 and p21 were increased. As for oxidative stress-inducing agents, no clear senescence-associated expression of genes was observed; only IL-8 levels significantly increased after Mena treatment in diffD9 and tBHQ treatment in diffD3 (Figure 6C and D).

**Figure 6:**
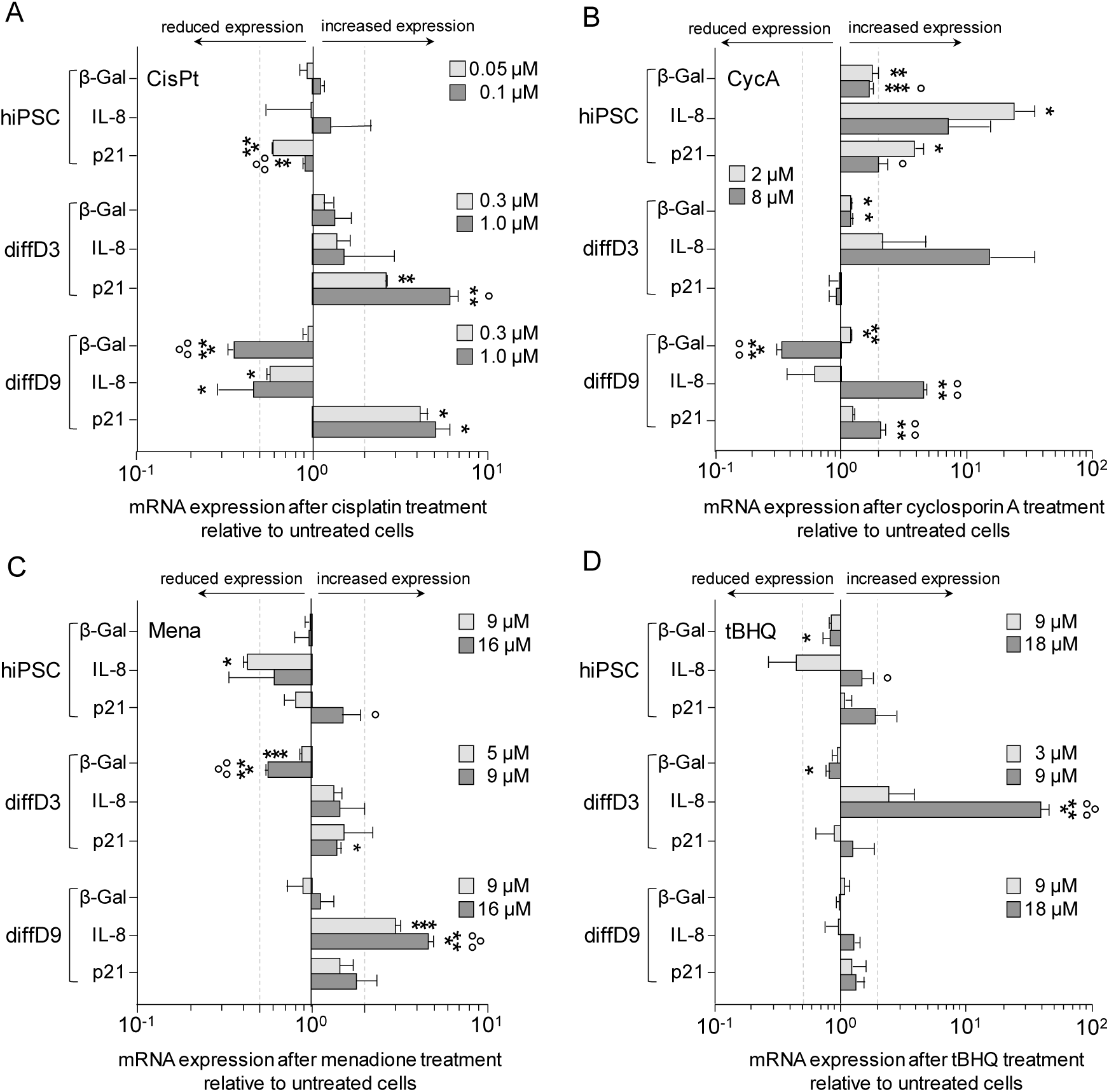
Influence of nephrotoxins and oxidants on the induction of senescence-related genes. Undifferentiated hiPSC, diffD3, and diffD9 were treated for 24 h with the indicated concentrations of CisPt (A), CycA (B), Mena (C) and tBHQ (D). 48 hours after treatment, quantitative RT-PCR analysis of the mRNA expression of senescence-associated genes (β-GAL, IL-8, p21) was performed. Data from at least three independent experiments are expressed as mean + SD. Student’s t-test, *p≤0.05 and **p<0.01 compared to untreated control One-way ANOVA, *p≤0.05, **p<0.01, ***p<0.001 versus untreated control, °p≤0.05, °°p<0.01, °°°p<0.001 versus cells treated with the lower toxin concentration. CisPt = cisplatin, CycA = cyclosporin A, diffD3/9 = differentiation day 3/9, β-Gal = β-Galactosidase, hiPSC = human induced pluripotent cells, IL-8 = Interleukin 8, Mena = menadione, p21 = cyclin-dependent kinase inhibitor 1, tBHQ = tert-butylhydroquinone.

### Effect of nephrotoxin and oxidant treatment during differentiation on marker expression and function of differentiated cells

The long-term impact of toxins on the differentiating cells was assessed by measuring marker expression and transport function in cells treated at diffD3 but evaluated at diffD9. As shown in Figure 7, all substances caused long term significant changes in the diffD9 cells. In general, it can be said that CisPt caused the most extensive changes, with all but two transcripts being significantly reduced. Surprisingly, the expression of the pluripotent stem cell markers Oct 3/4 and Nanog was also decreased. Oct 3/4 was used as a differentiation reference; the expression of the other genes was corrected by the amount of Oct 3/4 and therefore its expression was not statistically evaluated. No significant changes were caused by the other substances that would be considered biologically relevant, with the exception of Aqp-1 for all substances used, Meg for Mena and Cad-16 for tBHQ.

**Figure 7:**
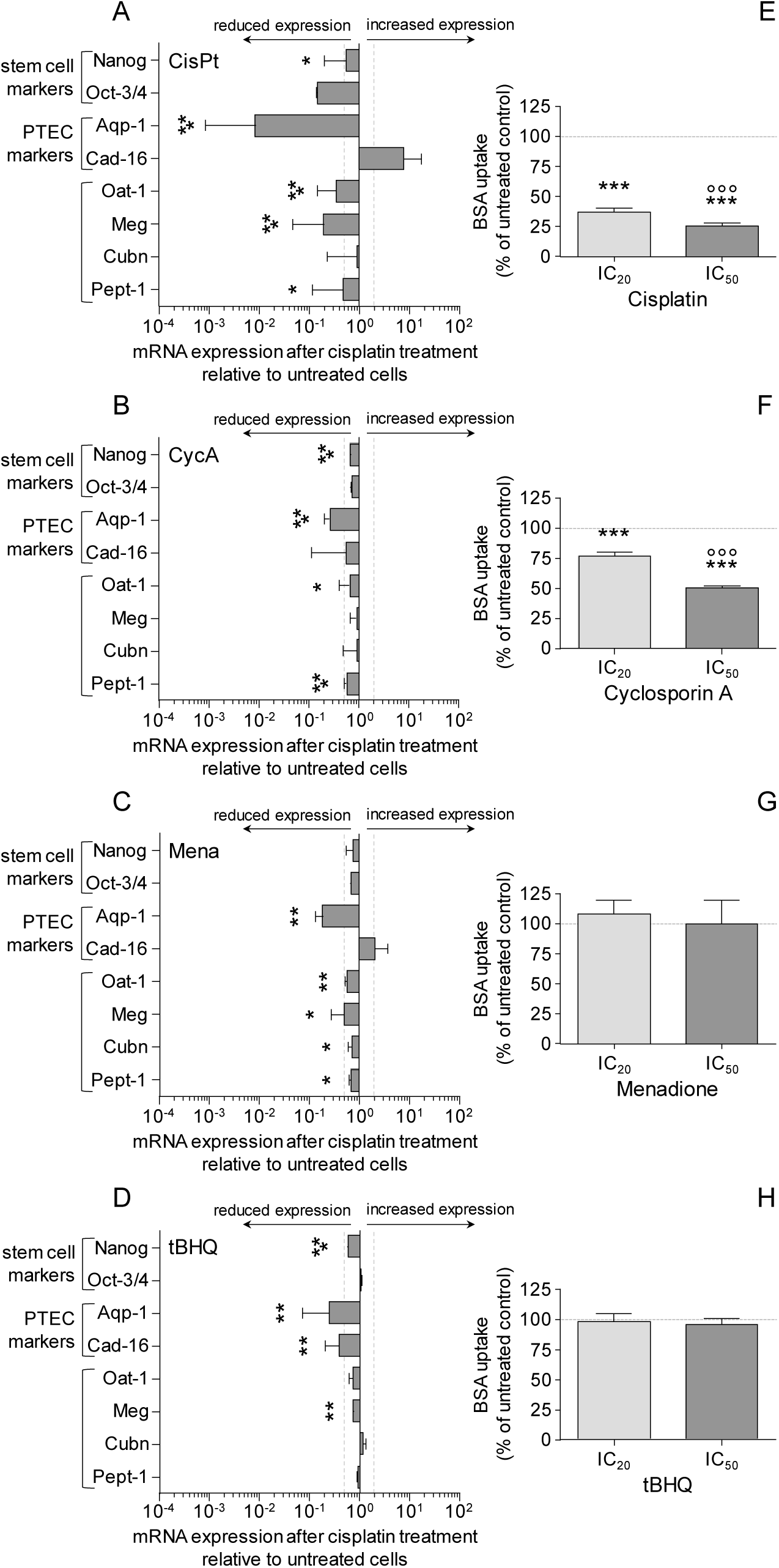
Influence of nephrotoxins and oxidants on marker gene expression and transport function during differentiation. Cells treated with the indicated concentrations of cisplatin and cyclosporin A on day 3 of the differentiation process, were analyzed on day 9 of differentiation and compared with untreated diffD9. Quantitative RT-PCR analysis of mRNA expression of diffD9 after 24 h of treatment with CisPt (A), CycA (B), Mena (C) and tBHQ (D) at diffD3 and analyzed on diffD9 for the mRNA expression of prototypic stem cell markers (Nanog, Oct-3/4), PTEC markers (Aqp-1, Cad-16) and transporters (Glut-5, Meg, Cubn, Pept-1, Oat-1), compared to their expression in untreated diffD9, set to 1. (E-H) Flow cytometric analysis of albumin uptake in diffD9 cells after 24 h of treatment with the hiPSC’s IC_20_ and IC_50_ of CisPt (E, 0.3 and 1.0 µM), CycA (F, 2.0 and 8.0 µM), Mena (G, 3.0 and 6.0 µM), and tBHQ (H, 4.0 and 8.0 µM) on diffD3 compared to untreated diffD9 using FITC-labelled albumin. Data from at least 3 independent experiments (exception: albumin uptake after IC_50_ treatment with tBHQ: 2 independent experiments) are expressed as mean + SD. *p≤0.05, **p<0.01, ***p<0.001 compared to untreated diffD9 (Student’s t-test or one-way ANOVA), °°°p≤0.001 vs. IC20 (one-way ANOVA). Aqp-1 = aquaporin-1, Cad-16 = cadherin 16, CisPt = cisplatin, Cubn = cubilin, CycA = cyclosporin A, diffD3/9 = differentiation day 3/9, hiPSC = human induced pluripotent cells, IC_20/50_ = inhibitory concentration where 20 and 50 % of the cells are dead, respectively, Meg = megalin, Mena = menadione, Pept-1 = peptide transporter 1, Oat-1 = organic anion transporter 1, Oct-3/4 = octamer-binding transcription factor 3/4, and Nanog = homeobox protein, tBHQ = tert-butylhydroquinone.

Treatment with cisplatin on differentiation day 3 had a strong effect on the cells’ ability to take up albumin on differentiation day 9. CycA also reduced this ability, but to a lesser extent. For both substances, the reduction in albumin uptake was dose-dependent. Mena had a low but significant effect on the expression of both parts of the albumin uptake complex, megalin and cubilin, but had no effect on albumin transport itself. tBHQ neither showed a relevant effect on the subunits of the albumin transport complex, megalin and cubilin nor on the uptake of albumin.

## Discussion

In this study, we explored the response of hiPSC to nephrotoxin (CisPt and CycA) and oxidant (Mena and tBHQ) exposure during and after their differentiation into PTELC. The obtained results demonstrate the high sensitivity of the differentiating cells to these agents. Depending on the agent, this sensitivity led to apoptosis or early signs of cellular senescence, which both have an unfavorable effect on the tubular regeneration process.

In a prior study, we successfully established a PTELC differentiation model from hiPSC using a protocol published in 2015 (Kandasamy et al. 2015), leading to hiPSC-derived PTELC with similar morphology to PTEC, expression of prototypical PTEC markers and the ability to undergo albumin endocytosis (Mboni-Johnston et al. 2023). When treated at different stages of PTELC differentiation with the nephrotoxins CisPt and CycA, in the case of CisPt, hiPSC and diffD3 were more sensitive to CisPt than differentiated PTELC, while all stages were equally sensitive to CycA [19] (Mboni-Johnston et al. 2023).

Here, we complemented the previous results with dose-response curves of the three differentiation stages, hiPSC, diffD3 and diffD9, after exposure to menadione and tBHQ to mimic oxidative stress occurring during kidney regeneration. Our findings revealed for the first time that diffD3 cells were more susceptible to these oxidants than undifferentiated hiPSC and fully differentiated PTELC. This observation is toxicologically relevant as it suggests that the cells differentiating into PTELC are particularly vulnerable to oxidative stress, which would be present during regeneration processes in an injured kidney (Gorin 2016). Consistent with this observation, our gene expression analysis revealed a decline in antioxidant defenses as cells differentiated, with essential antioxidant genes downregulated in diffD3 compared to diffD9, possibly explaining the sensitivity of the differentiating cells to oxidative stress. This was previously shown in human embryonic stem cells (Saretzki et al. 2008). The increased expression of redox associated genes in the differentiated PTELC compared to diffD3 may have contributed to their increased resistance against oxidative stress. Based on the low expression of redox genes in diffD3, we speculated that pluripotent stem cells differentiating into PTELC most likely have significantly reduced redox properties on the third day of differentiation. As a result, their antioxidant defenses would be weakened, and the damage induced by the oxidants would increasingly exceed the buffering capacity of the cell, leading to a significant reduction in viability.

The sensitivity of human iPSC and thereof differentiated counterparts to oxidizing agents has not been extensively investigated, resulting in a lack of readily available IC_50_ values in the literature. diffD9 demonstrated a significant reduction in cell viability by tBHQ at an IC_50_ of 35 μM, which is in the same range of endothelial cells and mammary epithelial cells with IC_50_ values of 60 μM (Karimi et al. 2019) and 50 μM (Jin et al. 2016), respectively. These results suggest that different cell types are sensitive to tBHQ to approximately the same extent. Regarding Mena, the human liver cell line HepG2 exhibited cytotoxicity to Mena with an IC_50_ value of approximately 18-19 µM after 18 h of incubation (Chen and Cederbaum 1997), which is very similar to the IC_50_ value of 17 µM observed in our differentiated cells after 24 h.

To uncover molecular mechanisms underlying the sensitivity of the various differentiation stages to the different toxins, proliferation, apoptosis and senescence were studied. The differing sensitivity of hiPSC and diffD3 towards CisPt compared to diffD9 may stem from variations in proliferation rates as indicated by the results obtained from the EdU incorporation measurements. This was expected to a certain extent, since CisPt is a DNA-damaging agent that has been reported to interfere with DNA synthesis and replication in rapidly dividing cells, such as stem cells (Wang and Lippard 2005). Two observations are interesting here: diffD3 cells have a proliferation rate that is about 50 % lower than that of hiPSC, but they are about as sensitive as hiPSC (Mboni-Johnston et al. 2023). It is therefore highly unlikely that the proliferation rate alone determines sensitivity. Otherwise, hiPSC would have to be even more sensitive than diffD3. Secondly, CisPt treatment did not significantly affect the proliferation rate of diffD3 compared to hiPSC, despite their similar sensitivity to CisPt. This suggests that the diffD3’s sensitivity to CisPt may be linked to other cytotoxic effects of the drug. Indeed, all studied stages of differentiation exhibited susceptibility to CisPt-induced cell death rather than senescence, as co-treatment with CisPt and the pan-caspase inhibitor QVD, a broad-spectrum caspase inhibitor that prevents apoptosis, effectively restored cell viability. This suggests a predominantly apoptotic cell death induced by CisPt. This result is somewhat unexpected as, in addition to apoptosis, CisPt is also recognized to induce other forms of cell death, such as ferroptosis and necrosis (Ikeda et al. 2021; Lee et al. 2001; Lieberthal et al. 1996). To the best of our knowledge, no study has investigated the mode of toxicity of CisPt in the tubular differentiation process yet. Only in pluripotent stem cells (Peskova et al. 2019) and fully differentiated PTEC (Kandasamy et al. 2015; Lawrence et al. 2022) the mode of toxicity of CisPt was examined up to now, with signs of apoptosis found in the stem cells and oxidative stress as well as DNA damage reported in the differentiated cells, in which apoptosis was not evaluated. Nevertheless, the results obtained in the present study are consistent with clinical data and the results of animal studies, revealing that CisPt exhibits dose-limiting nephrotoxicity and is apoptotic to renal proximal tubular epithelial cells of humans and experimental animals (Ciarimboli 2014; Miller et al. 2010).

When studying the mode of toxicity of the non-genotoxic nephrotoxin CycA at the different stages of differentiation, we found that the proliferation rate of hiPSC and diffD9 cells was particularly susceptible to CycA treatment. While the effect of CycA has not yet been studied in detail in pluripotent stem cells, the impact of CycA on the proliferation rate we observed is consistent with previous studies of renal cell types (Lally et al. 1999; Seki et al. 2005; Sun and Wang 1997). As the use of QVD revealed that CycA did not induce exclusively apoptotic cell death, we investigated whether the observed decrease in proliferation is related to the activation of possible senescence mechanisms. Indeed, we observed that CycA preferentially stimulated early signs of senescence in hiPSC but not in diffD3 and diffD9, which is in contrast to previous studies by Jennings and colleagues who showed an induction of cellular senescence in differentiated human PTEC (Jennings et al. 2007). However, since we only analyzed the mRNA expression of selected senescence-associated marker genes, further experiments are required for clarification. Nonetheless, the lack of evidence of early signs of senescence in diffD3 and diffD9 cells suggests that their sensitivity to CycA was not dependent on senescence activation. Consistent with this consideration, CycA was previously shown to induce apoptotic cell death mechanisms in a renal proximal tubular cell line (Healy et al. 1998; Lally et al. 1999).

Further studies on the molecular mechanisms contributing to the preferential hypersensitivity of differentiating cells to oxidizing agents showed that Mena and tBHQ inhibited proliferation and induced apoptosis only in differentiating cells. Comparable to our investigations with the two oxidants, Xiao and colleagues investigated the effect of H_2_O_2_-induced oxidative stress on bone marrow stem cells (MAPC), their differentiation into endothelial cells, and underlying mechanisms *in vitro* (Xiao et al. 2014). They could only observe significant inhibition of proliferation and induction of apoptosis in the MAPC and not in MAPC differentiating into endothelial cells (Xiao et al. 2014).

In addition to the effects of Mena and tBHQ treatment on cell viability at different stages of tubular differentiation, further analysis revealed that these oxidizing agents also selectively impaired the functionality of fully differentiated PTELC and the expression of individual marker genes associated with tubular differentiation. However, while Mena exposure significantly decreased mRNA expression of the PTEC transporter megalin, which could be an explanation for the loss of ability to take up albumin, tBHQ, on the other hand, increased megalin expression, leading to a corresponding increase in albumin uptake in differentiated PTELC. Kurosaki et al. showed that H_2_O_2_-induced oxidative stress like tBHQ also increased megalin expression in the HK-2 cell line (Kurosaki et al. 2018), suggesting similar effects of these two oxidants. Megalin is an important endocytic receptor that is highly expressed in PTEC in the kidney and is involved in the uptake of proteins, including albumin and certain drugs that are filtered in the glomeruli via the megalin/cubilin endocytosis system (Nielsen et al. 2016). Other toxic substances with oxidative stress-inducing properties, such as CisPt, are known to bind to megalin and prevent its interaction with its ligand, which includes albumin (Hori et al. 2017). To our knowledge, the impact of Mena or tBHQ on megalin has not yet been reported. Additional studies are required to elucidate how these oxidative stress-inducing agents affect megalin expression in PTEC. In addition to megalin, we found that both compounds significantly reduced the expression of the PTEC marker aquaporin-1 in diffD9, which has not yet been published. However, Liu et al. (Liu et al. 2018) showed that mitochondrial oxidative stress in obstructive kidney disease also led to the downregulation of aquaporins, including aquaporin-1.

Treatment during differentiation had a slight to strong effect on gene expression and albumin uptake, depending on the substance used. None of the substances upregulated the expression of the pluripotent stem cell markers, which implies that no substance generally prevented differentiation into PTELC. However, the significant reduction in mRNA expression of the key PTEC marker Aqp-1 indicates that all tested substances prevented the surviving population of diffD3 from differentiating efficiently into fully functional differentiated PTELC. In CisPt and CycA treated cells, this loss of functionality was apparent by the measured reduced albumin uptake capacity of diffD9, while no lower transport was observed in the oxidant-treated diffD9. Further functional tests are needed to assess the state of the diffD9. There are some reports of impairment of differentiation of various stem and progenitor cells (myogenic differentiation of myoblasts, osteogenic differentiation of mesenchymal stem cells, erythropoietic differentiation) by the accumulation of uremic toxins in chronic kidney disease, but no such reports for stem cells that differentiate into proximal tubule cells. (Alcalde-Estevez et al. 2021; Duangchan et al. 2022; Kamprom et al. 2021). But in immortalized renal progenitor cells a toxin, arsenite, also led to an altered differentiation, directing the differentiation into distal tubular cells, since a distal tubular marker protein, calbindin, was highly increased, while no significant change was detected in Aqp-1 (Singhal et al. 2023). So, our initial results and those from others suggest a potential vulnerability of the renal tubular epithelial regeneration process to nephrotoxic noxae, a finding that could potentially explain the observed loss of renal epithelial function after regeneration from AKI.

In conclusion, this study provides the first comprehensive characterization of the effects of the nephrotoxins CisPt and CycA, as well as the oxidizing agents Mena and tBHQ, on the tubular differentiation process in a 3R compliant model. The small selection of compounds analyzed here already underscores the diverse responses of cells at different stages of differentiation to substances with varying mechanisms of action. While it is apparent that the DNA-damaging compound CisPt had its most significant effect on rapidly growing proliferating cells, the deleterious effect of oxidants on differentiating cells was unexpected. Consequently, these observations lend support to our hypothesis that nephrotoxins and oxidative stress may impede tubular epithelial regeneration.

## Author contributions

N.S. and J.A. designed the study, I.M.M.-J., S.H. and C.K. performed research and analyzed the data, N.S., I.M.M.-J., S.H., J.A. and C.B.: drafted and revised the manuscript, all authors approved the final version of the manuscript.

## Ethical aproval

The manuscript does not contain animal studies, clinical studies or patient data.

## Acknowledgements

The outstanding technical assistance of Kerstin De Mezzo is acknowledged. This research was funded by the Deutsche Forschungsgemeinschaft (DFG, German Research Foundation) - 417677437/GRK2578.

