## Supplementary Table S1 for "Impact of nephrotoxins and oxidants on survival and transport function of hiPSC-derived renal proximal tubular cells"

**Supplementary Table S1. Primer sequences used for real-time PCR.**

ACTB = β-actin, AQP1 = aquaporin-1, CAD16 = cadherin 16, CAT = Catalase, CD13 = alanyl aminopeptidase, CUBN = cubilin, , β-GAL = beta galactosidase, GLUT5 = glucose transporter 5, GPX1 = glutathione peroxidase 1, GSS = glutathione synthetase, HMOX1 = Heme oxygenase 1, IL-8 = Interleukin 8, MEG = megalin, MnSOD2 = manganese-dependent superoxide dismutase, NANOG = homeobox protein, N-CAD = N-cadherin, NOS2/3 = Nitric oxide synthase 2/3, NOX1/4 = NADPH oxidase 1/4, NQO1, quinone oxidoreductase 1, NRF2 = Nuclear factor erythroid 2-related factor 2, OAT1/3 = organic anion transporter 1/3, OCT3/4 = octamer-binding transcription factor 3/4, p21 = cyclin-dependent kinase inhibitor 1, p47 = p47phox, a subunit of NADPH oxidase, PEPT1/2 = peptide transporter 1/2, RPL32 = ribosomal protein L32, SOD1 = superoxide dismutase 1, TRX = Thioredoxin, TXNRD1 = Thioredoxin reductase

| Gene | Forward | Reverse |
|---|---|---|
| **ACTB** | GAGCACAGAGCCTCGCC | TCATCATCCATGGTGAGCTGG |
| **AQP1** | CATCCTCTCAGGCATCACCTC | CACACCATCAGCCAGGTCATTG |
| **CAD16** | AGCACGTGTGAAGTCGAAGT | ACTGAGGTTCTGGGAAGTGATG |
| **CAT** | CAAAATGCTTCAGGGCCGC | GAGCACGGTAGGGACAGTTC |
| **CD13** | TGGCCACTACACAGATGCAG | CTGGGACCTTTGGGAAGCAT |
| **CUBN** | TAGCTTCGTGAAGGTGTGGG | GACTGGAAGACGGCAGTGAA |
| **β-GAL** | TGCGCAATGCCACCCA | CAGGGCACATACGTCTGGAT |
| **GCLC** | ACGGAGGAACAATGTCCGAG | CAGGACAGCCTAATCTGGGAA |
| **GLUT5** | GCCAAAGTGCACCCAGAATG | GTCAGCCTCCCTTCCTTCAT |
| **GPX1** | CCGGGACTACACCCAGATGA | TTGGCGTTCTCCTGATGCC |
| **GSS** | ATAGCTGCTGGCCGAAAACT | TCCGTGAGTCCCACTGTC |
| **HMOX1** | CTGCTCAACATCCAGCTCTTTG | CTTGGTGTCATGGGTCAGCA |
| **IL-8** | TTGGCAGCCTTCCTGATTTCT | GGGTGGAAAGGTTTGGAGTATG |
| **MEG** | GCCAGTGGCCAAGAATGTGA | TCCGCGTCATCTGAACAGTC |
| **MnSOD2** | GCTTTCTCGTCTTCAGCACC | AGATACCCCAAAACCGGAGC |

| | | |
|---|---|---|
| **NANOG** | ACCTCAGCTACAAACAGGTGAA | AAAGGCTGGGGTAGGTAGGT |
| **N-CAD** | AGGCTTCTGGTGAAATCGCA | GCAGTTGCTAAACTTCACATTGAG |
| **NOS2** | CTCCACATTGTTGTTGAT | AATCCAGATAAGTGACATAAG |
| **NOS3** | TGGAGTCTTGTGTAGGATA | CAAGGAGACGAAGAGAAC |
| **NOX1** | AATGTCACATACTCCACTG | CTCTCCAGCCTATCTCAT |
| **NOX 4** | TGACAGGTTTGTTGTCCTG | CTGGAAGAACCCAAGTTCCA |
| **NQO1** | ACCTTGTGATATTCCAGTTCCCC | GAACACTCGCTCAAACCAGC |
| **NRF2** | AGTGACTGAAACGTAGCCGA | CAGCTTTTGGCGCAGACATT |
| **OAT1** | AGTATGGAGGTACTCCGGGC | GCATGGAGAGGCAGAGGAAG |
| **OAT3** | CTTTGTGCCCTTGGACTTGC | GGAAGAGGCAGCTGAAGGAG |
| **OCT3/4** | ACCCACACTGCAGCAGATCA | CCACACTCGGACCACATCCT |
| **p21** | AGTCAGTTCCTTGTGGAGCC | GACATGGCGCCTCCTCTG |
| **p47** | GGGCGCGGATTTATAGCAGT | CCCCAGTCACTCACGTTTCC |
| **PEPT1** | CAAGTGCATCGGTTTTGCCA | CTCTTTAGCCCAGTCCAGCC |
| **PEPT2** | CTGGGAGGACAAGTGGTACA | AGTCCGTTCCTCTGCATGTT |
| **RPL32** | GTTACGACCCATCAGCCCTTG | CATGATGCCGAGAAGGAGATGG |
| **SOD1** | GCCTCATAATAAGTGCCATA | TCTGTTTCAATGACCTGTATT |
| **TRX** | GATGGTCAAGAGCCCAACCA | CCGGGAAGTATCTCGGTGTG |
| **TXNRD1** | AGCATGTCATGTGAGGACGG | CCAATTCCGAGAGCGTTCCT |
